# Surprise About Sensory Event Timing Drives Cortical Transients in the Beta Frequency Band

**DOI:** 10.1101/254060

**Authors:** T. Meindertsma, N.A. Kloosterman, A.K. Engel, E.J. Wagenmakers, T.H. Donner

## Abstract

Learning the statistical structure of the environment is crucial for adaptive behavior. Humans and non-human decision-makers seem to track such structure through a process of probabilistic inference, which enables predictions about behaviorally relevant events. Deviations from such predictions cause surprise, which in turn helps improve inference. Surprise about the timing of behaviorally relevant sensory events drives phasic responses of neuromodulatory brainstem systems, which project to the cerebral cortex. Here, we developed a computational model-based magnetoencephalography (MEG) approach for mapping the resulting cortical transients across space, time, and frequency, in the human brain (N=28, 17 female). We used a Bayesian ideal observer model to learn the statistics of the timing of changes in a simple visual detection task. This model yielded quantitative trial-by-trial estimates of temporal surprise. The model-based surprise variable predicted trial-by trial variations in reaction time more strongly than the externally observable interval timings alone. Trial-by-trial variations in surprise were negatively correlated with the power of cortical population activity measured with MEG. This surprise-related power suppression occurred transiently around the behavioral response, specifically in the beta frequency band. It peaked in parietal and prefrontal cortices, remote from the motor cortical suppression of beta power related to overt report (button press) of change detection. Our results indicate that surprise about sensory event timing transiently suppresses ongoing beta-band oscillations in association cortex. This transient suppression of frontal beta-band oscillations might reflect an active reset triggered by surprise, and is in line with the idea that beta-oscillations help maintain cognitive sets.

**Significance statement:** The brain continuously tracks the statistical structure of the environment to anticipate behaviorally relevant events. Deviations from such predictions cause surprise, which in turn drives neural activity in subcortical brain regions that project to the cerebral cortex. We used magnetoencephalography in humans to map out surprise-related modulations of cortical population activity across space, time, and frequency. Surprise was elicited by variable timing of visual stimulus changes requiring a behavioral response. Surprise was quantified by means of an ideal observer model. Surprise predicted behavior as well as a transient suppression of beta frequency band oscillations in frontal cortical regions. Our results are in line with conceptual accounts that have linked neural oscillations in the beta-band to the maintenance of cognitive sets.

## Introduction

Humans and other organisms continuously adapt their behavior to the statistical structure of their environment. This suggests that the brain is equipped with neural machinery for statistical learning, which can interact with the processes driving goal-directed behavior. Of particular importance here is surprise (Dayan and Yu, 2006; O’Reilly et al., 2013), a violation of one’s expectation about the next event, which might indicate a sudden change in the environmental structure, and can transiently boost central arousal state, increasing the organism’s sensitivity and learning rate (Yu and Dayan, 2005; Nassar et al., 2012).

Expectation, uncertainty, and surprise are intricately related. The precision of expectations scales with uncertainty, that is, the width of the distribution of observed events: high uncertainty precludes forming precise expectations. Violations of expectations cause surprise, the level of which depends on the difference between the expected and actually observed event (often termed prediction error). These intuitions can be formalized within the framework of Bayesian statistics and used to search for neurophysiological correlates (see Materials and Methods: *Bayesian ideal observer model: General approach and rationale*).

One important dimension of the environment is the timing of relevant sensory events (Gibbon et al., 1997; Nobre et al., 2007). Two lines of work have studied the neural basis of temporal expectation effects. One has shown that environments with rhythmic (i.e., precise) temporal structure entrain neural oscillations in the cerebral cortex, the phase of which then modulates sensory cortical responses, perception, and cognition (Lakatos et al., 2008; Schroeder and Lakatos, 2009; Rohenkohl and Nobre, 2011; Rohenkohl et al., 2012; Riecke et al., 2015; van Ede et al., 2017). In these rhythmic changes of the environment, surprise is minimized (once the structure is learned expectations match observations). Consequently, this first line of work has identified neural correlates of temporal expectation, rather than of surprise.

The second line of work has studied neural responses of subcortical, neuromodulatory centers, specifically, dopaminergic centers of the midbrain, to sensory events entailing reward. Because event timing here varied non-periodically from trial to trial as in many natural environments, this work could link phasic neuromodulatory responses to temporal surprise (Hollerman and Schultz, 1998; Fiorillo et al., 2008). Surprise-driven phasic responses might also occur in other neuromodulatory brainstem systems, such the noradrenergic system (Dayan and Yu, 2006). Because brainstem neuromodulatory systems have widespread projections to the cortical networks underlying goal-directed behavior, one would expect changes in cortical population activity elicited by surprise (Bouret and Sara, 2005). However, this second line of work on temporal expectation has focused on surprise-related activity in subcortical systems.

Here, we studied responses to surprise about the timing of sensory events in human cortex. A computational model-based magnetoencephalography (MEG) approach enabled us to map surprise-related cortical transients across space, time, and frequency. We used a Bayesian model that accumulated previously experienced durations of the interval between visual changes into posterior beliefs about the next interval duration. This ideal observer model provided trial-to-trial measures of temporal surprise, which predicted modulations of prefrontal and parietal cortical beta-band dynamics.

## Materials & Methods

This paper reports a reanalysis of an MEG data set that has previously been used for a study into decision-related feedback signals in visual cortex (Meindertsma et al., 2017). Here, we focus on those aspects of the experimental design that are most relevant for the issue addressed in the current paper: uncertainty and surprise about the timing of the experimental events specified below. We refer to our previous paper (Meindertsma et al., 2017) for a more detailed description of the visual stimulus and the behavioral task.

### Participants

Thirty-one volunteers participated in the experiment. Two participants were excluded due to incomplete data and one participant did not complete the experiment due to poor quality of simultaneously acquired pupil data. Thus, 28 participants (17 female, age range 20 - 54 years, mean age 28.3, SD 9.2) were included in the analysis. All participants had normal or corrected-to-normal vision and no known history of neurological disorders. The experiment was conducted in accordance with the Declaration of Helsinki and approved by the local ethics committee of the Hamburg Medical Association. Each participant gave written informed consent.

### Stimulus

MEG was measured while subjects viewed the intermittent presentation of a target (full contrast Gabor patch; diameter: 2°) and reported the on- and offset of the target (Figure 1A). Gabor targets flickered at 10 Hz (counter-phase) through alternation of two out-of phase Gabors every 50 ms. This caused steady-state evoked responses over visual cortex at 10 and 20 Hz (data not shown), distinct in terms of spectral profile, topography and functional characteristics from the surprise-related modulations we focused on here. The target was located in either the lower left or lower right visual field quadrant (eccentricity: 5°, counterbalanced between subjects), surrounded by a rotating mask (17°×17° grid of black crosses), and superimposed on a gray background. The mask rotated at a speed of 160°/s. The target was separated from the mask by a gray “protection zone” subtending about 2° around the target (Bonneh et al., 2001). Subjects fixated on a fixation mark (red outline, white inside, 0.8° width and length) centered on the mask in the middle of the screen. Stimuli were presented using the Presentation Software (NeuroBehavioral Systems, Albany, CA, USA, RRID:SCR_002521). Stimuli were back-projected on a transparent screen using a Sanyo PLC-XP51 projector with a resolution of 1024×768 pixels at 60 Hz. Subjects were seated 58 centimeters from the screen in a whole-head magnetoencephalography (MEG) scanner setup in a dimly lit room.

**Figure 1:**
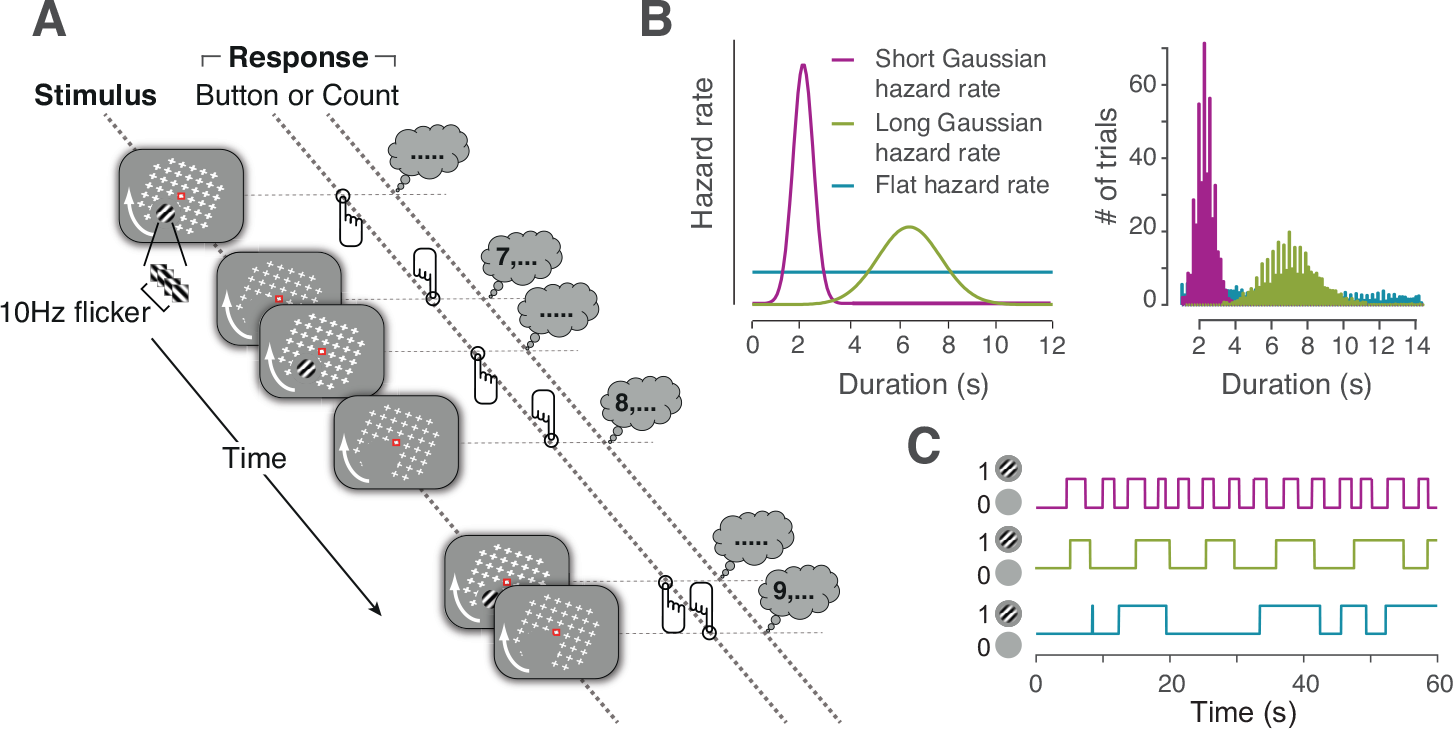
Behavioral task. **A.** Schematic depiction of the stimulus and task. A salient, flickering target (Gabor patch) temporarily appeared and disappeared on a rotating background. Subjects fixated on the red fixation mark and reported stimulus changes either by direct button press or silently counting the disappearances and reporting the total number at the end of the run. **B.** The interval duration between stimulus changes was randomly drawn from one of three distributions that corresponded to three hazard rates (left), resulting in distinct distributions of intervals (right, average histogram over subjects). **C.** Example time courses of target presence (1 = present, 0 = absent) drawn from these distributions.

### Behavioral task and experimental design

The subjects’ task was to maintain stable fixation and detect the physical offsets and onsets of the target, the predictability of which fluctuated from trial to trial, and the mean predictability of which varied systematically across blocks. To this end, the interval durations between stimulus changes were sampled from three different distributions in the different blocks. These distributions were computed so as to produce three predetermined so-called hazard functions, which describe the probability that an event will occur at a particular time, given that it has not occurred yet. The hazard function formalizes the expectation of a change and affects human reaction times in simple detection tasks (Luce, 1986). The hazard function can be computed as follows:

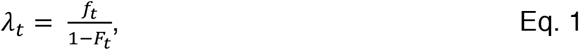

where *λ*_*t*_ is the value of the hazard function at time point *t*, *f*_*t*_ is the value of distribution *f* on time point *t*, and *F*_*t*_ is the area under the curve of distribution *f* from −*∞* to time *t*.

We used the following procedure to construct three ‘environments’, referred to as ‘Short’, ‘Long’, and ‘Flat’ below. We first selected three hazard functions that systematically differed in their level of predictability (Figure 1B, C). We then computed the actual distributions of intervals by re-arranging Eq. 1 as follows:

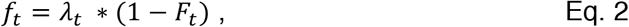

The interval durations were then randomly selected from *f*. Specifically, the temporal environments were defined as follows:

*Short:* The hazard function was a narrow Gaussian distribution with a mean of 2 s and a standard deviation of 0.2 s. This resulted in nearly periodic and, thus, largely predictable intervals between events.
*Long:* This condition used the same hazard function as the previous condition, but with a larger mean and standard deviation (6 s and 0.6 s, respectively) thus rendering event timings less predictable (Fiorillo et al., 2008).
*Flat:* The hazard function was flat with a mean of 6s, yielding the least predictable interval durations. The resulting distribution of interval durations, f_t_, therefore, approximated an exponential distribution; characterizing a memory-less process (i.e. the timing of the next event could not be predicted from previously encountered intervals, Feller, 1959).

Computational analysis with a Bayesian model (Fig. 2) described below confirmed that the sampled intervals from these three environments gave rise to different mean levels of uncertainty and surprise (Fig. 2G,H). The three environments were presented in separate three-minute blocks.

Within each of the above temporal environments, there were two behavioral tasks. Both tasks required subjects to monitor the changes of the small visual target. In one task (called *Detection-button*), they were asked to report those changes immediately. Specifically, subjects reported target offsets or onsets by pressing a button with the right index or middle finger, respectively. In the other task (*Detection-count*), they were asked to count and report the changes at the end of the block. Subjects silently counted the number of target offsets and reported the total in response to a 4-AFC question at the end to the block. The two tasks were randomly selected before each block under the constraint that both would occur equally often. We here analyzed both task conditions, but only found robust effect for Detection-button.

All subjects completed a total of 6 blocks of the Short environment, and 16 blocks of the other Long and Flat environments, resulting in about the same number of trials per environment. Additionally, subjects performed a motion-induced blindness task and a functional localizer task, which were not relevant for the current study, but are reported in our previous paper (Meindertsma et al., 2017). All blocks within an environment were completed in succession; the order of environments was counter-balanced across subjects.

### Bayesian ideal observer model: General approach and rationale

We developed an ideal observer model to quantify surprise and uncertainty about the timing of sensory events (i.e., the target on- and offsets). The model tracked the evolving predictive distribution of upcoming interval durations; more specifically, it computes the posterior predictive of unobserved interval durations, conditional on the observed data, throughout each block of the experiment. We assumed that subjects tracked the temporal statistics of the task in a similar way, and we used the posterior predictive distribution as a proxy of the subjects’ belief states (i.e., their prediction of the timing of the next stimulus change).

While we used an ideal observer model that prescribed the optimal inference for our task, we are agnostic to the precise inference process that was used by our subjects and we do not claim that subjects used the exact computations used by the model. Our central assumption was that subjects accumulated observations throughout each block (i.e., over more than just one or two previous intervals). This assumption was derived from a substantial body of work on other forms of learning and evidence accumulation (Sutton and Barto, 1998; Gold and Shadlen, 2007; Glaze et al., 2015), and it was supported by the findings described in Results. Our model implemented the normative accumulation strategy by perfectly integrating across the entire history of the observations (here: of interval durations) and updating internal representations accordingly. A practical benefit of this approach was that it did not require fitting of model parameters, for which our current data did not provide sufficiently strong constraints.

The only free parameter in the model was the level of temporal estimation noise, which we allowed to scale with the magnitude of the interval duration according to Weber’s law (Gibbon et al., 1997). To this end, we transformed the discrete values of the observed intervals into Gaussian distributions that were used to update the model (see next section). The mean of these distributions was equal to the observed interval *t* and their standard deviation was equal to the observed interval *t* times a Weber’s fraction (coefficient of variation, Gibbon et al., 1997). We simulated the model with 34 Weber’s fraction values ranging from 0.001 to 0.5 (0.001, 0.05:0.01:0.35, 0.4, 0.5). We then computed the correlation between the measured single-trial reaction times (pooled across all subjects) and surprise (see *Bayesian ideal observer model: Implementation*), separately for each Weber fraction, and selected the Weber fraction that maximized this correlation. To this end, we fitted a second order polynomial to the correlation coefficients as a function of Weber’s fraction and extracted the maximum of the polynomial. This yielded a Weber’s fraction of 0.17 (Figure 2F), which was used for all analyses reported in this paper. Using model-based surprise from a noise-free version of the model yielded qualitatively identical results (data now shown).

### Bayesian ideal observer model: Implementation

We assumed that the subjects used a model in which the observed intervals have been generated from a gamma distribution with parameters alpha (shape) and beta (scale). These parameters were given uninformative prior distributions (Lee and Wagenmakers, 2013), which were updated by the data to posterior distributions.

Using the interval duration distributions as the observations, we could obtain the expectations about to-be-observed intervals by generating posterior predictives (i.e., drawing an alpha-beta pair from the joint posterior distribution and then drawing a predicted interval from the associated gamma distribution; repeating this process many times yields a posterior predictive distribution for the to-be-observed interval). We assumed that the subjects updated their belief state after each observation of a new interval duration. Likewise, the model was updated after every interval *t* by computing a new posterior predictive distribution, based on the durations of intervals *1:t* and the prior.

We generated a posterior predictive distribution over the to-be-observed intervals using Gibbs sampling (a Markov chain Monte Carlo, or MCMC, algorithm; Andrieu et al., 2003) in the software JAGS (Plummer, 2003) and Matlab (version R2013a, RRID:SCR_001622). We used two Markov chains with different starting points comprised of 2500 samples per chain with 500 samples burn-in, for a combined total of 4000 samples. The posterior predictive MCMC samples Y_1…4000_ for the next interval, *t+1*, were then summarized by a gamma distribution using the functions ‘gamfit’ and ‘gampdf’ in Matlab (Figure 2A,B):

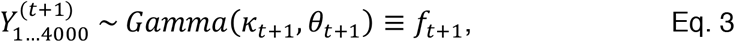

where 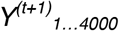 are the MCMC samples, and *κ*_*t+1*_ and *θ*_*t+1*_ are the parameters of the gamma distribution *f*_*t+1*_; hence, *f*_*t+1*_ is the continuous posterior predictive distribution for the upcoming interval after having observed the preceding intervals *1…t*.

To be able to relate trial-to-trial uncertainty and surprise to behavior and the MEG data, we extracted two information theoretic metrics from the time-evolving posterior predictive distribution *f*_*t+1*_ (i.e., belief).

*Uncertainty:* We quantified trial-to-trial uncertainty about the timing of the upcoming interval *t+1* as the entropy of the posterior predictive distribution f_t+1_ (i.e., the posterior predictive based on intervals *1…t*):

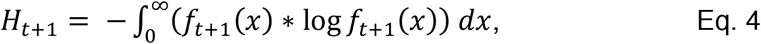

where *H*_*t+1*_ is the entropy after intervals *1*…*t*, and the integral is over all possible values *x* for the upcoming interval. Entropy depended on the width of *f*_*t+1*_, and thus uncertainty was higher when predictions of interval durations were less precise (Figure 2A,C,D). For clarity, in what follows we will use the term entropy when referring to this uncertainty.

*Surprise:* For every upcoming interval *t+1*, we computed the surprise about the corresponding interval duration in terms of the Shannon information conveyed by the interval duration *x*_*t+1*_, given the posterior predictive distribution *f*_*t+1*_:

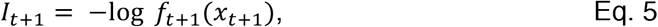

where *I*_*t+1*_ is the information gained by interval *t+1*, given *f*_*t+1*_. Thus, surprise was defined as the negative log-probability of the upcoming interval *t+1*, given the intervals that had been presented so far.

We added one further transformation in the computation of surprise. The surprise measure defined in Eq. 5 quantified the surprise about the upcoming event timing based on the posterior predictive distribution *f*_*t+1*_, but disregarding the time elapsed in the current interval. It is unlikely that exactly this distribution translated into subjects’ level of surprise: as time passed and no event occurred in a given interval, all interval durations shorter than the elapsed time become impossible. Subjects likely discounted these impossible intervals in their expectation of the timing of the upcoming event, which should have also affected their level of surprise. In other words, their internal representation of the posterior predictive distribution changed dynamically throughout each trial, as a function of elapsed time. We constructed a time-varying version of the posterior predictive distribution *f*_*t+1*_, which was also conditioned on the elapsed time on interval *t*. This version was equal to *f*_*t+1*_ for elapsed time equal to 0 and then increasingly deviated from *f*_*t+1*_ as elapsed time grew. We approximated this time-varying distribution, denoted as *f*’_*t+1*_ in the following, by setting all probabilities in *f*_*t+1*_ up to the current time point to zero and renormalizing the remaining distribution to integrate to 1 (Figure 2B). We then computed surprise based on this new distribution *f*’_*t+1*_ using Eq. 5. The time-variant prior *f*’_*t+1*_ converged to 1 as time passed, and thus surprise approached zero for longer intervals.

### Regressing computational variables against behavior

We used reaction time (RT) during Detection-button as behavioral readout of the impact of uncertainty and surprise. Accuracy approached ceiling for all subjects, due to the high saliency of the target. We computed and compared mean RTs per environment and stimulus event (target off- and onset).

We also correlated RT to our trial-to-trial estimates of surprise and entropy. RT was log-transformed so as to normalize the skewed RT distributions. To test if the model-based surprise fitted the behavioral (RT) data better than a linear combination of just the two previous interval durations (i.e., a leaky accumulation with strong leak), we used multiple linear regressions to compare the following two nested models:

M1: *log(RT) ~ Interval_t_*Env*Event + Interval_t-1_*Env*Event*
M2: *log(RT) ~ Interval_t_*Env*Event + Interval_t-1_*Env*Event + Surprise*Con*Event*,

where *Interval_t_* and *Interval_t-1_* corresponded to the durations of the two intervals preceding the visual change (i.e., interval on trial *t* and *t-1*), and *Surprise* was the computational model-derived metric. Predictors were multiplied by categorical variables envrionment (*Env*, the three different temporal environments) and *Event* (target offset or onset). Both variables strongly affected RT (Figure 3A). We fitted both M1 and M2 and compared the fits per subject using adjusted R^2^.

### MEG data collection

Magnetoencephalography (MEG) data were acquired on a CTF 275 MEG system (VSM/CTF Systems, Port Coquitlam, British Columbia, Canada) with a sample rate of 1200 Hz. The location of the subjects’ head was measured in real-time using three fiducial markers placed in the both ears and on the nasal bridge to control for excessive movement. Furthermore, electrooculogram (EOG) and electrocardiogram (ECG) were recorded to aid artifact rejection. All data were recorded in sets of four blocks of three minutes duration (or two blocks at the end of an environment set).

### MEG data analysis

#### Preprocessing

The data were analyzed in Matlab (version R2013a, The Mathworks, Natick, MA, USA, RRID:SCR_001622) using the Fieldtrip (Oostenveld et al., 2011, RRID:SCR_004849) toolbox and custom-made software.

#### Trial extraction

In blocks involving subjects’ reports, we extracted trials of variable duration, centered on subjects’ button presses, from the 3 min blocks of continuous stimulation. We call this method for trial extraction “response-locked”. The following constraints were used to avoid mixing data segments from different percepts when averaging across trials: (i) The maximum trial duration ranged from −1.5 s to 1.5 s relative to report; (ii) when another report occurred within this interval, the trial was terminated 0.5 s from this report; (iii) when two reports succeeded one another within 0.5 s, no trial was defined; (iv) for the analysis of Detection-button blocks, we included only those reports that were preceded by a physical change of the target stimulus within 0.2 to 1 s, thus discarding reports not related to stimulus changes. We used this method for the analyses related to surprise. In an alternative analysis of all Detection blocks, trials were defined in the same way as described above, but now aligned to physical target on- and offsets (“stimulus-locked”). In the Detection-count task, no button responses were given during the block, so stimulus-locked trial extraction was the only option. We used this method for the analysis related to entropy (see Kloosterman et al., 2015b & Meindertsma et al., 2017 for a similar procedure).

#### Artifact rejection

All epochs that contained artifacts caused by environmental noise, eye-blinks, muscle activity or squid jumps were excluded from further analysis using standard automatic methods included in the Fieldtrip toolbox. Epochs that were marked as containing an artifact were discarded after every artifact detection step. For all artifact detection steps the artifact thresholds were set individually for all subjects. Both of these choices aimed at optimization of artifact exclusion. Line-noise was filtered out by subtracting the 50, 100, 150 and 200 Hz frequency components from the signal.

#### Time-frequency decomposition

We used a sliding window Fourier transform to compute the time-frequency representation for each sensor and each trial of the MEG data. The sliding window had a length of 200 ms and a time step size of 50 ms, with one Hanning taper (frequency range 5-35 Hz, frequency resolution 5 Hz and frequency step size 1 Hz). The data was baseline corrected for every frequency bin and MEG sensor separately. The baseline was computed by averaging single-trial power over a baseline time window. The baseline time windows ranged from −1.25 to −0.75 s for response-locked and −1 to −0.5 s for stimulus-locked analyses, respectively. The time course of every frequency bin and sensor combination was baseline corrected by subtracting the single-trial baseline power at that frequency and dividing by the mean baseline power across trials within an experimental environment. We used the single-trial baseline power for subtraction to eliminate the effect of slow power fluctuations, because any surprise-related power modulations could only have occurred after the sensory event that elicited surprise. We used the mean baseline for division in order to minimize noise in the single-trial estimates of the single-trial power modulation values. This division was used to compensate for the common decay of power with frequency, which hinder identification of effects at higher frequencies, and to normalize the single-trial modulation values (Siegel and Donner, 2010). It did not systematically alter the association of power modulation values with other variables.

#### Source reconstruction

We used an adaptive linear spatial filtering method called linear beamforming (Van Veen et al., 1997; Gross et al., 2001) to estimate single-trial modulations of MEG power at the source level. We computed a common filter for a baseline time window (1 to 0.5 s before response), a ‘transient’ time window, and a frequency band of interest (0 to 0.5 s after response, 20 Hz +/− 4 Hz spectral smoothing, see dashed box in Figure 4A). The transient time window and frequency band of interest were selected based on cluster-based statistics at the sensor level (see next section). We used the measured head positions and individual single-shell volume conductor models, based on individual images from T1-weighted structural MRI. We computed the power values, in both baseline and transient time windows, for each trial and source grid point (i.e., voxel) as follows. First, we projected the sensor-level MEG power values from the time window of interest as well as from a baseline time window through the common spatial filter. Second, we converted the estimated power values during the time window of interest into units of power modulation, again by subtracting and dividing by the corresponding baseline power values.

### Correlating single-trial computational variables to MEG power

We correlated the MEG power modulation to our measures of entropy and surprise, as derived using our model (see *Bayesian ideal observer model: Implementation*) across trials. Although intricately related (see Introduction), uncertainty and surprise entailed different computations (see above). A key difference was *when* during the course of a trial these two quantities where computed. So, we reasoned that neural correlates of these computational quantities should also differ in their dynamics: uncertainty about event timing should be reflected in the neural baseline state before occurrence of the sensory event, whereas surprise should be reflected in a transient response elicited by that event. Thus, we used different components of the single-trial MEG power estimates for the analyses of entropy and surprise.

#### Entropy

We correlated entropy to the MEG power modulation separately in every MEG sensor and frequency bin. This was done within subject and separately for the three environments. There are structural differences in entropy and surprise between these environments (Figure 2G,H), thus pooling over these conditions might result in inflated correlations that reflect session differences instead of the true correlation between entropy and MEG power. We reasoned that entropy should affect baseline or tonic arousal, where high entropy should cause higher arousal. As our task was continuous, we considered the time window right before the stimulus change the best reflection of a baseline state. For this reason we averaged the MEG power over the time period right before a stimulus change (−0.5 to −0.25s with respect to the target offset or onset) before correlating to entropy.

The results were then averaged over the three environments and transformed with the Fisher z transformation (Fisher, 1915):

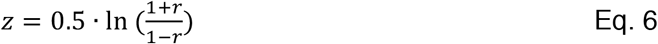

We used two-tailed permutation tests with a cluster-based correction for multiple corrections to test the correlation coefficients against zero (Efron and Tibshirani, 1998; Maris and Oostenveld, 2007).

#### Surprise

Correlations between surprise and MEG power modulation were performed using the same method, with the following exceptions. First, we attuned the analysis in two ways to account for the correlation between surprise and RT (Figure 2F, 3). Because of this correlation, any post-stimulus correlations between surprise and MEG power modulation might reflect differences in the timing of the button press. We performed this analysis response-locked, because these RT differences are difficult to disentangle from genuine effects of surprise when the power modulations are time-locked to the stimulus change. Additionally, to account for confounding effects of RT and the duration of the previous interval, we also performed a partial correlation analysis between surprise and MEG power modulation with the interval duration preceding the stimulus change or RT as covariate. Second, for the correlation between surprise and MEG power modulation we did not average over a specific time window, but instead performed correlations separately for every time point, resulting in a 3-dimensional matrix of correlations (sensor * frequency bin * time point). Consequently, we also performed cluster-based permutation statistics over these three dimensions. The correlations that survived cluster correction were visualized by integrating (i.e. computing the area under the curve) over sensors and frequency bins (for the time course), sensors and time points (for the frequency spectrum), frequency bins and time points (for the topography) or just over sensors for the time frequency representation (see Hipp et al., 2012 for a similar approach).

To assess the robustness of the emerging clusters we performed a cross-validation analysis using a leave-one-out procedure. To this end, we repeated the cluster-based permutation statistics on all possible iterations of N-1 subjects, each time using the resulting cluster as a mask to calculate the average correlation in the left-out subject, separately for target offset and onset trials. These values were tested against zero and against each other across subjects using permutation tests (10.000 permutations).

We also computed the correlation between trial-to-trial power modulation averaged over the whole cluster and log(RT). The resulting correlations were tested against zero across subjects using a permutation test (10.000 permutations).

The transient modulations of MEG power estimated for each voxel in the source grid, derived by means of source reconstruction (see *MEG data analysis: Source reconstruction*), were correlated to the trial-to-trial measure of surprise. This was done separately within each subject and the resulting correlations averaged over subjects after Fischer’s z-transformation (Eq. 6). For comparison, we also computed the average modulations of MEG power in the same time window and frequency band. The resulting maps of correlation or average power modulation were nonlinearly aligned to a template brain (Montreal Neurological Institute) using the individual images from structural MRI. To test the similarity of the spatial topography of the correlation to the average modulation of power, we correlated the two corresponding source maps per subject and tested the correlation coefficients again zero on the group level by means of a permutation test (10.000 permutations).

## Results

Subjects (N=28) performed a simple visual detection task reporting on- and offsets of a small, but salient target stimulus (Figure 1A). In different blocks, target events were administered using three different temporal environments (Figure 1B,C) translating into different overall levels of uncertainty and surprise about the timing of target events (Figure 2G,H). In order to quantify these two computational variables not only across conditions, but also across individual trials, we developed a Bayesian belief-updating model. The model incorporated the evolving beliefs (i.e. the posterior predictive distributions) of an ideal observer about the temporal intervals between the sensory events. Beliefs were dynamically updated across trials and even within trials (for surprise, see *Materials and Methods*). From these time-evolving probability distributions, we extracted trial-by-trial measures of information-theoretic entropy (quantifying uncertainty) and surprise, which we related to the behavior and neural dynamics of our participants.

**Figure 2:**
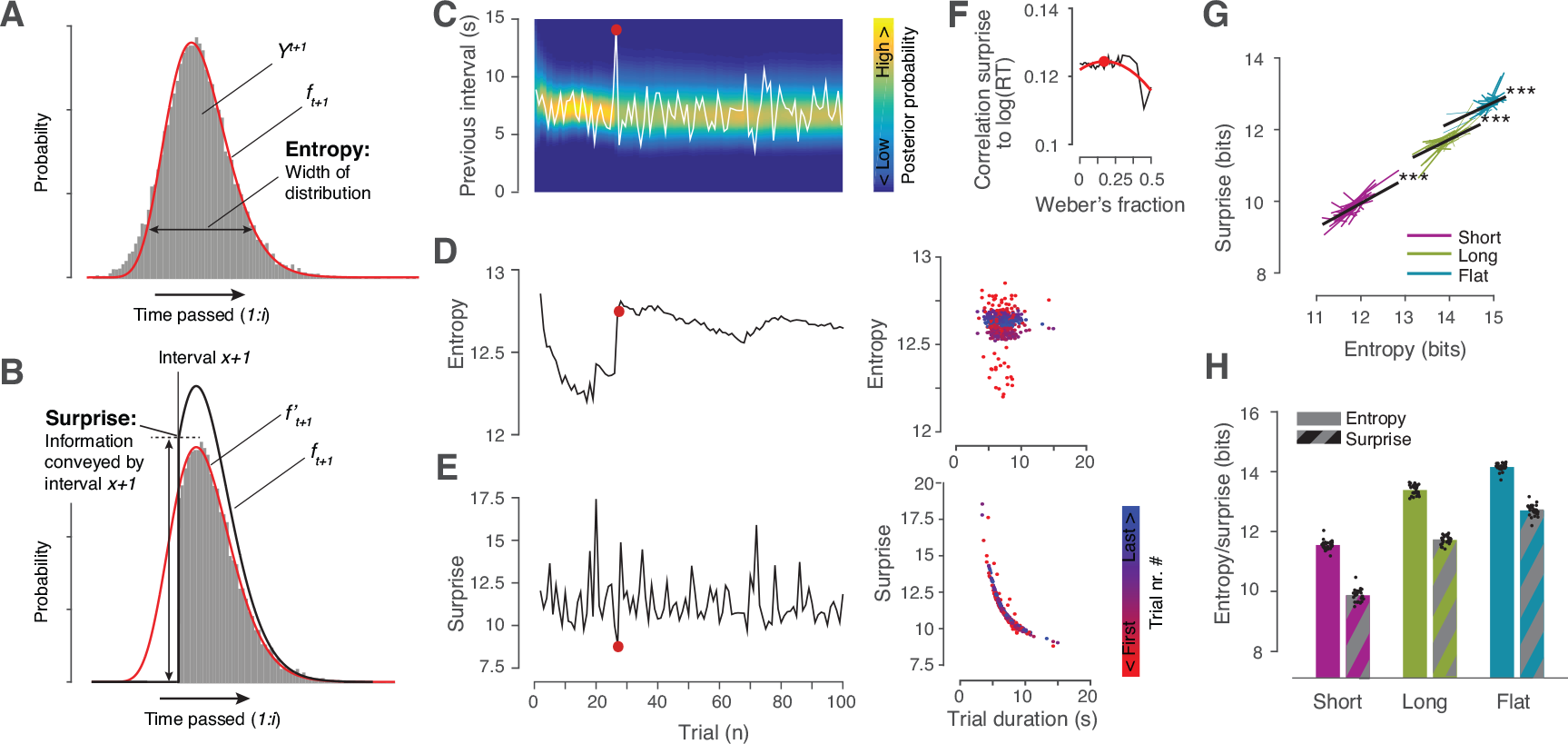
Bayesian updating model of belief about temporal structure. **A-F.**The model estimated the posterior predictive distribution over timings of stimulus changes for each upcoming interval *t+1*. This distribution is denoted as *f*_*t+1*_. The gray histogram shows the distribution of MCMC-samples (*Y^t+1^*) from the posterior predictive distribution for interval *t+1*. *f*_*t*_ was estimated by fitting a gamma probability density function (red line) to *Y*^*t+1*^; it was then used to extract two different information-theoretic computational variables for each trial: entropy and surprise. **A.** Entropy, a measure of the uncertainty about the timing of the interval duration of the current interval, computed from the complete distribution *f*_*t+1*_ using Eq. 4 (see main text). The wider the distribution, the higher entropy. **B.** Surprise, a measure of information provided by each new interval duration, was also computed from the posterior predictive distribution, but with one extra step (see main text): the part of the distribution up to the current interval duration was truncated, and the remainder of the distribution re-normalized to integrate to 1 (*f*’_*t+1*_, black line). Surprise was defined based on this truncated function using Eq. 5 (see main text). **C.** Example sequence of interval durations (white line, from the long Gaussian condition) with posterior predictive distribution *f* (color coded). **D.** Entropy corresponding to interval durations in panel C (left); relationship between interval duration and entropy (right). **E.** Surprise, analogous to panel D. Red dot: example of exceptionally long interval (see duration in panel C). Surprise on this trial was low (panel E) because time dependent surprise decreased over time. After observing this interval entropy increased (panel D) because the observed interval was longer than the expected duration, given previous intervals. **F.** Correlation between log(RT) and surprise as a function of different Weber fractions (black line, see Materials and Methods). Second-order polynomial fit over these correlations used to select the Weber fraction yielding peak correlation. (red line; red dot depicts peak = 0.17). **G.** Regression of surprise on entropy. Thin colored lines, regression lines of single subjects; black lines, group average regression. **H.** Trial-averaged surprise and entropy for the three experimental environments defined in Figure 1. Bars, group average; black dots, single subjects. *** p = 0 for all tests, permutation tests across subjects.

Estimates of entropy and surprise fluctuated across trials, especially in the early part of each block (Figure 2C-E). The trial averages of both measures within each block also varied lawfully between the different experimental conditions, scaling with the predictability of the stimulus changes (Figure 2G,H). Estimates were smallest for the Short condition, intermediate for the Long condition, and largest for the Flat condition. As expected, variations in entropy and surprise were weakly correlated across trials (r=0.13 for Short and Flat, r=0.19 for Long condition, Figure 2G), since both measures were computed from the same probability distribution (Materials and Methods). Even so, these two variables entailed distinct computations, possibly by distinct neural circuits. Critically, both computational variables could be computed at different times during a trial, thus possibly leading to different dynamical modulations cortical population activity.

### Surprise predicts reaction time

The model-derived computational variable surprise predicted subjects’ reaction time (RT) in the detection task. Mean RT scaled with the different temporal environments in the same way as surprise and entropy, with the fastest RTs for Short and slowest RTs for the Flat condition (Figure 3A, compare with Figure 2E). RT correlated with surprise also at the single trial level (Figure 3B). We did not find robust correlations to RT for entropy.

We also tested whether model-derived surprise (entailing accumulation of intervals across the entire experimental block) predicted RT over and above a linear combination of only the two previous intervals (entailing, e.g. a leaky accumulation with strong leak). To this end, we used a nested regression model, which quantified the predictive power of a combination of surprise and the previous two intervals in accounting for the influence of temporal environments and target on- or offset, on RT (Materials and Methods). We compared this against a simpler model with only the two previous intervals. Because model-based surprise depended on all previous intervals, the comparison between the above two nested models assessed the impact on reaction time of intervals beyond the second one. We used adjusted R^2^ for comparison, which penalized model complexity. This comparison yielded higher adjusted R^2^ values for the model including surprise in 22 of 28 subjects (Figure 3C), indicating that surprise predicted RT over and above the duration of the previous two intervals.

Taken together, these results indicate that subjects tracked the temporal structure of the task by accumulating interval distributions at least over more than two intervals, akin to what was prescribed by the ideal observer model. We next searched the whole-brain MEG data for a dynamical neurophysiological signature of this process. To this end, we focused on the trial-to-trial fluctuations of surprise within each of the environments (Short, Long, Flat), which were more pronounced than the differences in mean surprise between environments.

**Figure 3:**
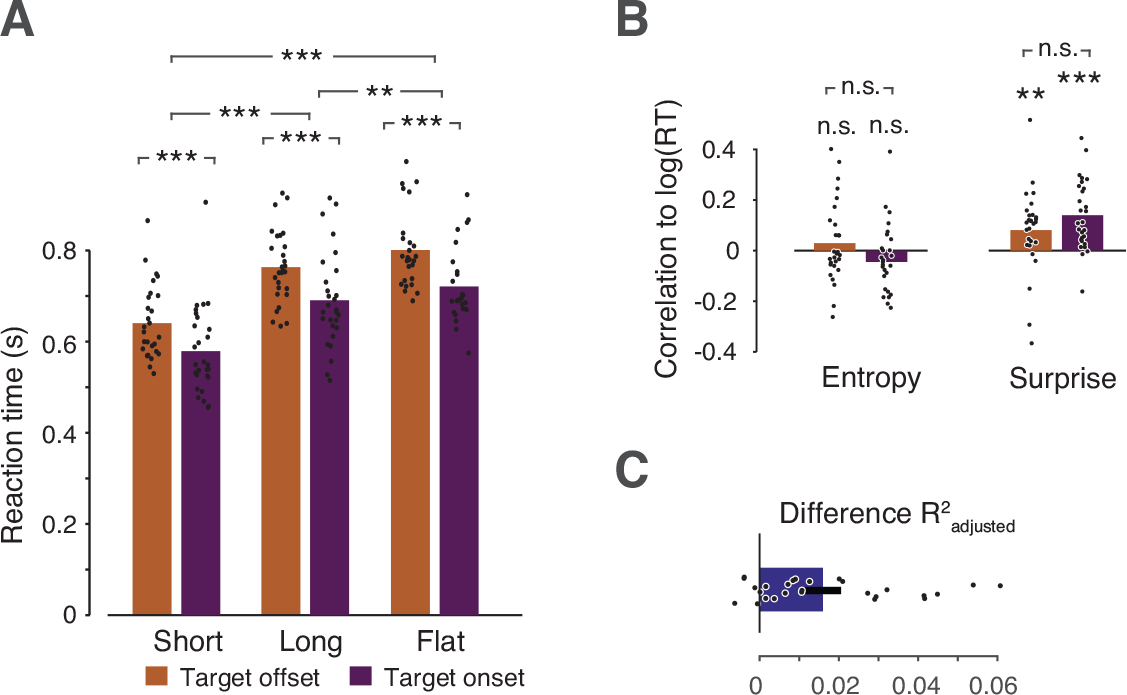
Link between computational variables and behavior. **A.** Average reaction time (RT) per interval distribution, separate for reports of target offsets and onsets. Bars show average over subjects; black dots depict average per subject. *** p<0.001, ** p<0.01, permutation tests across subjects, 10.000 permutations (differences between target off- and onsets, p-values were p=0, p=0 and p=0.003, for Short, Long and Flat, respectively; between conditions, p=0, p=0 and p= 0.001, for Short-Long, Short-Flat and Long-flat, respectively). **B.** Correlation between trial-to-trial fluctuations in Entropy (left)/Surprise (right) and log-transformed RT. Mean correlation coefficient Entropy: mean r=0.03 (s.d.=0.02), p=0.17 and r=-0.04 (s.d.=0.03), p=0.07 for off- and onset, respectively, difference off-on p=0.47; Surprise: mean r=0.09 (s.d.=0.02), p=0.007 and r=0.15 (s.d.=0.03), p=0 for off and onset, respectively, difference off-on p=0.48. **C.** Model comparisons between partial (including only 2 previous intervals) and full regression models (further including surprise; see main text). Difference in adjusted R^2^. Positive values indicate support for full model. Dots depict individual subjects; bars depict the mean, error bars depict SEM.

### Widespread cortical beta-band transient driven by surprise

We mapped out the cortical responses to trial-to-trial fluctuations in surprise by correlating the model-based surprise measures to modulations of MEG power, around the time of subjects’ behavioral responses to sensory events. We did this in an exhaustive fashion across every time and frequency bin and MEG sensor and tested for clusters of significant correlations across these three dimensions, while applying cluster-based multiple comparison correction (*Materials and Methods*). This approach revealed negative correlations in the beta (~20 Hz) frequency range, as well as in the lowest frequency bin resolved (5 Hz), indicating that higher surprise was associated with lower power in these frequency ranges. The peak in this negative correlation cluster started about 0.2 s before and reached its maximum about 0.25 s after subjects’ report of the stimulus change. This cluster exhibited several peaks over central, left frontal, and to a lesser extent left parietal cortex (Figure 4A,C).

For all analyses shown in Figures 4 and 5, we used partial correlations, controlling for reaction time, and we focused on the Detection-button task that entailed immediate behavioral report of the change of the visual target (Materials and Methods). We controlled for reaction time because (i) the data showed that the latter was affected by surprise (Figure 3), and (ii) motor responses are known to modulate beta-power around the time of response (Donner et al., 2009). Thus, button-presses could have potentially influenced the modulation by surprise. We focused on Detection-button because (i) we could only establish links between surprise and behavior for this task and (ii) it allowed us to lock neural dynamics more closely to the conscious registration of the visual change. When performing the correlation analysis for the Detection-count task (then locked to the physical stimulus change), we did not obtain any in significant correlation clusters. In two control analyses, we confirmed that the above results were robust to (i) using the ‘raw’ correlation between surprise and MEG power and (ii) controlling for the preceding interval duration. Both analyses resulted in highly similar clusters of negative correlations (data not shown).

The surprise-related cluster was robust and not driven by outliers, and the effect was not specific to the type of stimulus event (target on- or offset). We used a leave-one-out cross-validation procedure to test the robustness of the correlations on both target on- and offsets (Materials and Methods). We found robust negative correlations in the left-out subjects (Figure 4D). Furthermore, the correlation was found for both target offsets and onsets (Figure 4D, mean −0.036, −0.025, SEM 0.010, 0.006, p = 0.006, 0.002, for target offsets and onsets, respectively; difference: p=0.26, permutation tests, 10.000 permutations).

As expected from previous work on modulations of MEG power around motor responses (Donner et al., 2009), the overall modulation of MEG power in the time-frequency window of the surprise-correlation cluster (16-24 Hz, 0–0.5 s from response, normalized by the baseline 1-0.5 s before response) peaked in bilateral motor cortex (Figure 4B). But the component of beta-power modulations that correlated with trial-by-trial surprise showed a different cortical distribution, with negative correlations that peaked in the central sulcus, extending from motor- to more frontal cortex, and in left frontal and parietal cortex (compare Figure 4B and 4C). Indeed, there was no similarity between the individual topographies of the surprise-linked and the overall power modulations (mean correlation across subjects: r=-0.02, p=0.42). These observations indicate that the report-locked modulation linked to surprise and of overall power were located in distinct cortical networks.

**Figure 4:**
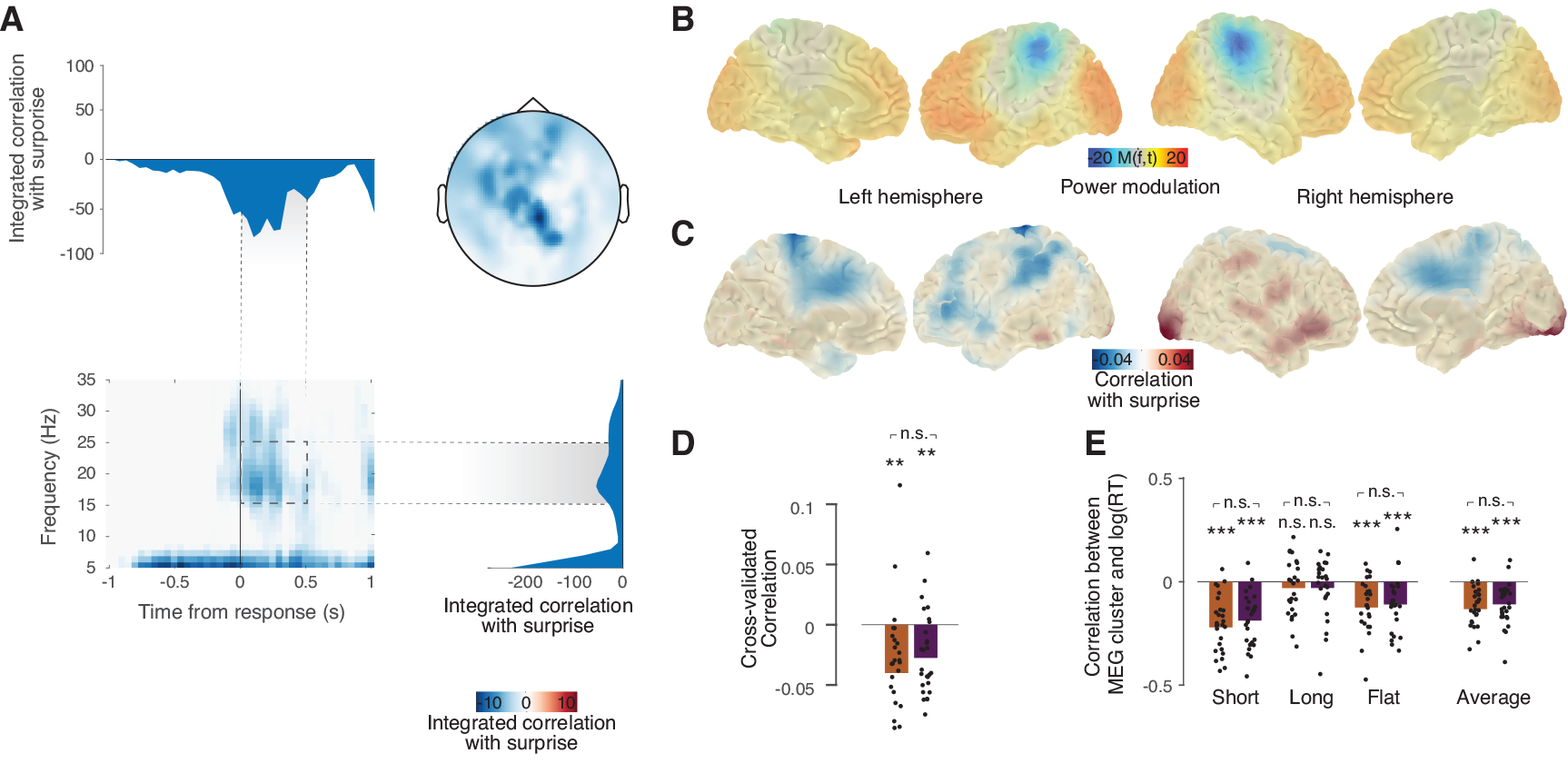
Widespread cortical beta-band transient driven by surprise. **A.** Exhaustive partial correlation (controlling for RT) between trial-to-trial measures of surprise and MEG power modulation in all sensors, time and frequency bins results in one cluster (cluster-based correction for multiple comparison, p = 0.001, two-sided) of negative correlation. Different panels show different dimensions of the cluster by integrating over the other dimensions; top left: time course, top right: spatial topography, bottom left: time-frequency representation; bottom right: frequency spectrum. **B.** Source reconstruction of the power modulation in the time window in which surprise-MEG correlation was strongest (dashed box in panel A). **C.** Source-reconstructed illustration of the correlation between transient modulation and trial-to-trial surprise depicted in panel B. These source maps are not statistically thresholded, but instead serve for comparing the correlation’s spatial distribution with the transient power modulation in panel B (average correlation between surprise and power modulation across subjects = 0.02, p=0.42). **D.** Leave-one-out cross-validation of the cluster found in panel A, separately for target offsets and onsets. Cluster-based permutation was performed on N-1 subjects and the average correlation in the resulting cluster was computed for the remaining subject (black dots); bars show averages over subjects. Correlation values were tested against 0 (permutation test; ** p<0.01; p=0.005 and p=0.002 for target offsets and onsets, respectively, p=0.19 for offsets-onsets difference). **E.** Correlation between MEG power in the cluster and log(RT) for separate distributions and average RT; permutation tests. All *** p <0.001, all offset-onset differences p>0.05 (lowest was p=0.21).

The surprise-related cluster for target offsets and onsets both exhibited a bimodal in the frequency domain, similar to the pooled analysis (Figure 5; compare to Figure 4A): next to the peak around 20 Hz just after response, an additional peak was evident in the lowest frequency bin resolved (5 Hz). For offsets, the effect was quite sustained in time (−0.25 to 0.5s around response); the topography showed peaks over parietal and occipital cortex and over left frontal cortex (Figure 5A). By contrast, the cluster for target onsets was more confined in time (with a sharp peak ~0.1s after report) and a different topography that peaked over central parietal cortex (Figure 5B). Taken together, our results suggest that perceptual surprise about both target on- and offsets elicited cortical transients in the beta-band. We consider them general dynamical correlates of temporal surprise monitoring. In addition, stimulus changes seem to have recruited additional processes expressed in the very low (<= 5 Hz) frequency range.

Finally, we asked whether the trial-to-trial fluctuations in beta-power modulations also predicted trial-to-trial variations in subjects’ (log-transformed) RTs. Here, we used the Pearson correlation values (i.e., without regressing out RT; *Materials and Methods*). Just as surprise, beta-power in the cluster also robustly predicted RT (Figure 4E). These correlations were negative, as expected based on the negative correlation between surprise and MEG-power (Figure 4A). We also compared the strength of this correlation between MEG-power and RT to the strength of the correlation between surprise and RT across, this correlation between correlations was positive, but not significant (r=0.19, p=0.33).

### No robust correlations between MEG baseline power and entropy

We did not find any evidence for a correlation of the raw baseline (−0.5 to 0 s with respect to stimulus change) MEG power with uncertainty, as measured in entropy. Correlations between entropy and MEG power spectra in the time window before stimulus change did not result in any significant (sensor-frequency) clusters that survived multiple-comparison correction (data not shown). It is likely that this lack of robust correlation reflected the continuous reduction in trial-to-trial variations of entropy over the course of each block (Figure 2C).

**Figure 5:**
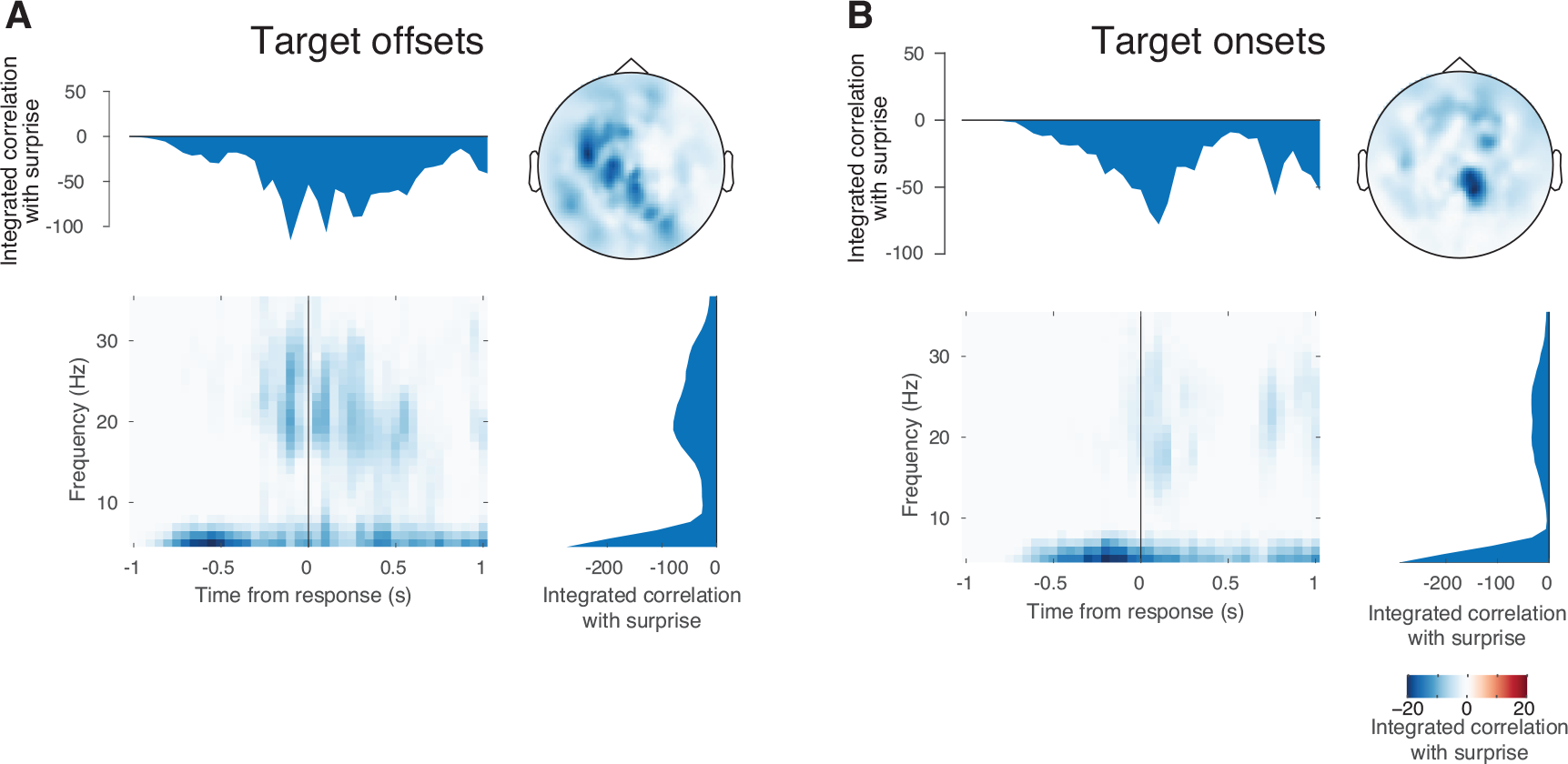
Separate correlations for offsets and onsets. Correlation analysis and cluster-based statistics performed separately for target offsets (**A**) and target onsets (**B**); p=0.001 for both analyses.

## Discussion

In this study, we comprehensively mapped cortical transients elicited by surprise about the timing of sensory events. We used a Bayesian updating model to estimate trial-to-trial variations of surprise and correlated these to subjects’ behavior as well as to neural dynamics, across the cortical surface. The model-derived surprise estimates predicted across-trial and environment variations in RT. The surprise estimates also predicted transient suppressions of low-frequency and beta-band power in a widespread network comprising motor-, prefrontal and parietal cortical regions, predominantly in the left hemisphere. The model-derived surprise estimates were more closely related to both behavior and cortical dynamics than the mere trial-to-trial variations in externally observable interval timings.

The signatures of surprise we uncovered in the beta frequency band were quite similar around target on- and offset (Figure 5). This stands in sharp contrast to the opposite beta-band modulation during (illusory or veridical) target disappearances and reappearances, proposed to reflect a decision-related feedback signal to in visual cortex (Meindertsma et al., 2017). The beta-band transients identified here likely reflected a distinct process that did not encode the content of the perceptual change, but rather the level of surprise about it.

One possibility is that surprise is computed in those fronto-parietal cortical networks exhibiting the surprise-related modulation of beta-oscillations observed here. Another possibility is that the surprise-related modulations are inherited from other regions projecting to those fronto-parietal networks. Indeed, neuromodulatory brainstem systems are a prominent candidate source. In particular the dopaminergic and noradrenergic systems are driven by temporal expectation and surprise (Aston-Jones and Cohen, 2005; Dayan and Yu, 2006; Fiorillo et al., 2008). Further, there is mounting evidence for a link between neuromodulation and beta-band power in visual cortex (Belitski et al., 2008; Donner and Siegel, 2011; Safaai et al., 2015; Zaldivar et al., 2018).

Specifically, phasic responses in dopaminergic nuclei, encode not only reward, but also the expected timing of reward arrival. The strength of these phasic neuronal responses inversely scales with predictability of the timing of reward, in line with encoding surprise about reward arrival, and it also predicted behavioral anticipation of reward (i.e. licking behavior) in monkeys (Fiorillo et al., 2008). Our current study complements this previous work, by unraveling the cortex-wide dynamics elicited by surprising events. Our design did not involve rewards but rather neutral, yet behaviorally relevant sensory events.

In a previous report based on the same data set as the current one (Kloosterman et al., 2015a), we showed that mean pupil dilation responses during the perceptual changes scaled in amplitude across the three environments in line with mean surprise as shown in the current Figure 2H. Pupil dilation is closely linked to phasic responses in neuromodulatory brain systems, in particular the noradrenergic locus coeruleus (Joshi et al., 2016; Reimer et al., 2016; de Gee et al., 2017). Thus, if the surprise-related modulations of cortical activity observed here were driven by phasic neuromodulation, one would expect to find correlations between single-trial pupil responses and surprise (Preuschoff et al., 2011; Nassar et al., 2012). Due to the sluggish dynamics of the peripheral pupil apparatus (Hoeks and Levelt, 1993; De Gee et al., 2014), testing for trial-by-trial correlations between pupil dilations and surprise (or, likewise, between baseline pupil diameter and uncertainty) in our experiment requires dedicated analysis approaches that tease apart fluctuating baseline levels and responses evoked by individual events. Using a general linear model (Hoeks and Levelt, 1993; De Gee et al., 2014), we failed to obtain reliable single-trial pupil responses and correlations to single-trial surprise (data not shown). This failure was likely, at least in part, due to the rapid nature of the current experimental design. Future work should use more widely spaced intervals to test whether pupil dilations reflect trial-to-trial variations of surprise.

Our current study provides a comprehensive picture of the cortical transients elicited by surprise, by systematically mapping these transients across the cortical surface and time-frequency plane. Previous work in humans has also studied neural correlates of model-derived measures of surprise, although this entailed surprise about stimulus identity, and not timing. Electrophysiological work found surprise about cue identity to modulate the P3 component of the EEG event-related potential as well as motor cortical excitability (Bestmann et al., 2008; Mars et al., 2008). Functional magnetic resonance imaging work linked surprise about the spatial location of stimuli to transients in posterior parietal cortex (O’Reilly et al., 2013). An EEG study dissociated oscillatory neural signatures of surprise and evidence accumulation (Gould et al., 2012). This latter study also found surprise-related modulation of beta-band power primarily at frontal and parieto-occipital electrodes, but the underlying cortical distribution was not estimated. Future studies of surprise in other domains (e.g. about cue identity) should use a similar approach to assess if surprise-related cortical transients are domain-general or -specific. Further, simultaneous EEG and MEG recordings (Schurger et al., 2015) are necessary to unravel the relationship between surprise-linked modulations of fronto-parietal beta-band oscillations and of the P3-component.

Another line of work has investigated the functional role of externally entrained low-frequency oscillations in temporal expectation. For fixed intervals, alpha phase in sensory cortices was found to be predictive of expected time of target arrival and lowered the threshold for sensory detection (Lakatos et al., 2008; Cravo et al., 2011, 2013; Rohenkohl and Nobre, 2011). Alpha oscillations might reflect rhythmic fluctuations in cortical excitability, entrained by rhythmic sensory input, which aids stimulus processing and perceptual performance (Schroeder and Lakatos, 2009). The high variability in interval durations (see Figure 1B,C inset) might explain the lack of alpha-band effects in our study. First, the range of possible durations was too broad to form predictions that fall within a specific phase of an alpha cycle. Second, even when oscillatory phase was modulated by temporal expectation in our task, the trial-to-trial variability would make it difficult to align trials and make these modulations visible.

It is tempting to relate our results to conceptual accounts of the functional role of beta-band oscillations in the brain (Engel and Fries, 2010; Spitzer and Haegens, 2017). One account (Engel and Fries, 2010) holds that beta-band oscillations help maintain the current sensorimotor or cognitive state (termed the ‘status quo’). Another account (Spitzer and Haegens, 2017) holds that beta-band oscillations help activate the currently relevant task sets. In both frameworks, the need for maintaining the current status quo, or task set, is low when surprise (the violation of expectation, or probability of change in the environment) is high, in line with our observation of a suppression of beta-band oscillations under high surprise.

While our current work presents an important first step towards unraveling the modulation of cortical dynamics by surprise, it is limited in that we only studied environments with constant statistical structure within each block. Once a posterior distribution has been learned, there remains no unexpected uncertainty, only expected uncertainty (Yu and Dayan, 2005). By contrast, the statistical structure of natural environments is often volatile. Richer experimental designs, that are volatile and include unmarked changes, allow for probing into richer, presumably hierarchical dynamics (Sugrue et al., 2004; Nassar et al., 2012; Meyniel et al., 2015). A more volatile task-environment would also lead to an increase in trial-to-trial variability of our entropy measure, providing a better-suited context to study the effects of this type of uncertainty on cortical processing. Our ongoing work aims to push beyond these limits by using richer environmental statistics that require more complex inference processes.

To conclude, we here uncovered a novel signature of temporal surprise that affected an elementary perceptual decision (target detection) and was characterized by a temporally focal, but spatially widespread, modulation of cortical population activity. This modulation might be instrumental in translating inferences about the behaviorally-relevant temporal structure into its consequences for action.

**Author contributions:**
Conceptualization, T.M., N.A.K., E.J.W. and T.H.D.; Investigation, T.M.; Formal analysis T.M.; Analytic tools, E.J.W. and N.A.K.; Writing - Original draft, T.M. and T.H.D; Writing - Review & Editing, T.M., N.A.K., A.K.E., E.J.W. and T.H.D.; Funding Acquisition, A.K.E. and T.H.D.; Supervision, T.H.D.

## Acknowledgements

This work was supported by the Netherlands Organization for Scientific Research (NWO, dossiernummer 406-14-016, to T.H.D. and T.M.); the Amsterdam Brain and Cognition priority program (ABC2014-01, to T.H.D.); the European Union Seventh Framework Programme (FP7/2007-2013) under grant agreement no. 604102 (Human Brain Project) (to T.H.D. and A.K.E.); and the German Research Foundation (DFG): Heisenberg Professorship DO 1240/3-1 (to T.H.D.), the Collaborative Research Centers SFB 936 (Projects A2/A3, A7, to A.K.E., T.H.D.) and TRR 169 (Project B1 to A.K.E.). We thank Udo Böhm for methodological advice and all members of the Donner lab for helpful discussion.

